# Slowing of Frontocentral Beta Oscillations in Atypical Parkinsonism

**DOI:** 10.1101/2022.04.19.488835

**Authors:** Marius Krösche, Silja Kannenberg, Markus Butz, Christian J. Hartmann, Esther Florin, Alfons Schnitzler, Jan Hirschmann

**Affiliations:** Institute of Clinical Neuroscience and Medical Psychology, Medical Faculty, Heinrich Heine University, 40225 Düsseldorf, Germany; Center for Movement Disorders and Neuromodulation, Department of Neurology, Medical Faculty, Heinrich Heine University, 40225 Düsseldorf, Germany

## Abstract

Diagnosis of atypical parkinsonian syndromes mostly relies on clinical presentation as well as structural and molecular brain imaging. It has not been investigated whether cortical oscillatory activity exhibits features distinguishing these syndromes. Therefore, we measured resting-state magnetoencephalography in 13 patients with corticobasal syndrome, 10 patients with progressive supranuclear palsy, 23 patients with idiopathic Parkinson’s disease and 23 healthy controls. We compared spectral power as well as amplitude and frequency of power peaks between the groups.

Atypical Parkinsonism was associated with spectral slowing, distinguishing both corticobasal syndrome and progressive supranuclear palsy from age-matched healthy controls and Parkinson’s disease. Patients with atypical Parkinsonism showed a shift of beta peaks (13-30 Hz) towards lower frequencies in frontal and central areas bilaterally. A concomitant increase in theta/alpha power relative to controls was observed in both atypical parkinsonian syndromes and in Parkinson’s disease.

Our results demonstrate that slowing of frontocentral beta oscillations is characteristic of atypical Parkinsonism. Spectral slowing with a different topography has previously been observed in other neurodegenerative disorders, such as Alzheimer’s disease, suggesting that spectral slowing might be an electrophysiological marker of cortical neurodegeneration. As such, it might support differential diagnosis of parkinsonian syndromes in the future.

**Highlights:**

- Slowing of beta oscillations distinguishes atypical parkinsonian syndromes from healthy controls and idiopathic Parkinson’s disease
- Spectral slowing is most pronounced in frontocentral areas
- Beta oscillations are shifted towards lower frequencies independent of their amplitude

## Introduction

Parkinsonism is a clinical syndrome characterized by various motor symptoms such as rigidity, tremor, bradykinesia, and abnormal gait (Lang and Lozano 1998). Most patients with Parkinsonism suffer from idiopathic Parkinson’s disease (PD) characterized by responsiveness to levodopa treatment and slow disease progression. PD is caused by neuronal loss in the substantia (Lang and Lozano 1998).

Atypical parkinsonian syndromes (APS) such as corticobasal syndrome (CBS) or progressive supranuclear palsy (PSP) resemble PD clinically. Yet, they have distinct pathologies and include a wide range of additional cognitive and motor symptoms (Burrell et al. 2014; Armstrong et al. 2013; Höglinger et al. 2017). The clinical hallmarks of CBS are asymmetric presentation of parkinsonism, apraxia and cognitive decline (Armstrong et al. 2013). Patients with PSP present with vertical supranuclear gaze palsy and postural instability (Höglinger et al. 2017; Litvan et al. 1996). Compared to PD, APS show a more rapid disease progression and poor responsiveness to levodopa (Armstrong et al. 2013; Litvan et al. 1996; Poewe and Wenning 2002).

Both CBS and PSP are tauopathies, characterized by aggregation of hyperphosphorylated tau protein in diverse brain regions, in conjunction with neuronal degeneration (Dickson et al. 2002; Kouri et al. 2011; Williams and Lees 2009). In this respect, they have commonalities with Alzheimer’s disease (Jack et al. 2013). Corticobasal degeneration (CBD), the pathology underlying CBS, is related to widespread tau aggregation in cortical areas and the basal ganglia (Dickson et al. 2002; Kouri et al. 2011; Forman et al. 2002), whereas the pathology of PSP is characterized by pronounced tau aggregation in the brainstem, with less cortical involvement (Dickson et al. 2007; Williams and Lees 2009; Dickson et al. 2010).

The relationship between pathology and symptomology is ambiguous for both CBS and PSP. Their clinical presentations are heterogeneous and overlapping (Williams and Lees 2009; Armstrong et al. 2013; Höglinger et al. 2017). Although diagnostic accuracy improves with disease progression and manifestation of symptoms, misdiagnosis is common (Osaki et al. 2004; Armstrong et al. 2013; Joutsa et al. 2014; Alexander et al. 2014). This does not only concern diseases within the APS spectrum but extends to other neurodegenerative diseases like PD (Hughes et al. 2002; Beach and Adler 2018) and AD (Di Stefano et al. 2016; Sakae et al. 2019). Since clinical examination alone is often not sufficient for an early and reliable differential diagnosis, it is of clinical relevance to identify new biomarkers.

Biomarkers might emerge from several sources, including imaging (Whitwell et al. 2010; Albrecht et al. 2017; Matsuda et al. 2020; Josephs et al. 2008; Dutt et al. 2016; Brenneis et al. 2004) and electrophysiology. The electrophysiology of APS, however, is not well investigated. Initial studies point to disease-specific alterations (Tashiro et al. 2006; Barcelon et al. 2019; Montplaisir et al. 1997), but a quantitative analysis of neuronal oscillations in APS and differences to other parkinsonian syndromes is missing. Here, we performed spectral analysis of resting-state magnetoencephalography (MEG) measurements in 13 CBS patients, 10 PSP patients, 23 PD patients and 23 age-matched healthy controls (HC). We describe for the first time a characteristic spectral signature of APS.

## Materials and methods

### Participants

For this study, we analyzed resting-state MEG measurements of 25 APS patients, 3 PD patients, and 11 HC. Two APS patients were excluded from analysis due to excessive noise. Of the remaining 23 patients, 4 were classified as possible CBS, 9 as probable CBS, 1 as possible PSP and 9 as probable PSP (Tab. 1). The diagnosis was made by an experienced movement disorders specialist (CH) based on current diagnostic criteria for CBS (Armstrong et al. 2013) and PSP (Litvan et al. 1996; Höglinger et al. 2017).

**Tab. 1:** Summary data of the study cohorts. CBS: Corticobasal Syndrome, PSP: Progressive Supranuclear Palsy, PD: Parkinson’s disease; HC: Healthy controls.

| Group | N | Mean age (years) | Min -Max age (years) | Gender (f/m) | Diagnosis (possible/probable) | Mean UPDRS-III (sum) |
| --- | --- | --- | --- | --- | --- | --- |
| CBS | 13 | 64 | 52 - 76 | 9 / 4 | 4 / 9 | 51.85 |
| PSP | 10 | 71.5 | 64 - 79 | 5 / 5 | 1 / 9 | 36 |
| PD | 23 | 63.73 | 47 - 79 | 4 / 19 | - | 36.32 |
| HC | 23 | 65 | 56 - 74 | 12 / 11 | - | - |

Additionally, we included unpublished resting-state MEG data of 20 PD patients of the akinetic-rigid or intermediate subtype and 12 HC. PD patients withdrew from dopaminergic medication over night before participation. For the CBS patients, Parkinsonism was quantified by the Unified Parkinson’s Disease Rating Scale III (UPDRS-III Goetz et al. 2008), cognitive abilities by the Montreal Cognitive Assessment (MoCA; Nasreddine et al. 2005), and apraxia by Goldenberg’s Apraxia Test (Goldenberg 1996) and the Test of Upper Limb Apraxia (TULIA; Vanbellingen et al. 2010). UPDRS-III scores were available for CBS and PD (except one) but not for PSP patients (Tab. S1). In a few cases, physical disabilities made it infeasible to measure some test items reliably. Thus, test scores were normalized by the number of scored items before computing correlations.

The groups significantly differed in age (One-way ANOVA: *F* = 3.15, *p* = 0.031). Post-hoc, two-sided t-tests revealed that the PSP group was significantly older than the CBS group (*p* = 0.018), the HC group (*p* = 0.005) and the PD group (*p* = 0.007). Excluding all PSP patients > 70 years eliminated the age difference and yielded similar results. The UPDRS-III scores did not significantly differ between CBS and PD (two-sided t-test: *p* = 0.081). All participants gave their written informed consent before participation. The study was approved by the local ethics committee in accordance with the Declaration of Helsinki (study numbers: CBS: 2019-447-andere; PSP: 2018-155-KFogU; PD: 5608R).

### Recordings

MEG recordings were conducted with a 306-channel whole-head MEG system (MEGIN, Espoo, Finland) in a magnetically shielded room. Participants were recorded for 10 minutes in upright position during rest with open eyes. A subset of 9 PD patients was measured for 5 minutes. Participants were requested to sit still, relax, and to not think of something specific. Electromyograms were recorded from both forearms to monitor upper limb movements.

### Data preprocessing

MEG data was preprocessed in MATLAB R2018a with the Fieldtrip Toolbox (version: 20201214; Oostenveld et al. 2011). Bad channels and time segments containing artifacts were discarded from further analysis based on protocol notes and visual inspection of sensor time series data, power spectra and topographies. Periods with tremor or other movements were discarded. After removing artifacts, gradiometer time series were down-sampled to 1000 Hz, 1 Hz high-pass filtered (FIR-filter) and subjected to an independent component analysis (Bell and Sejnowski 1995). Components containing eye blink or heartbeat artifacts were removed from the continuous gradiometer time courses based on their time series properties and topography.

### Source & Parcel Reconstruction

MEG data and individual T1-weighted MRI scans (Siemens Magnetom Trio, 3T MRI scanner) were co-registered and used to construct forward models via the single shell approximation method (Nolte 2003). In one case (PD12), the colin27 template (Holmes et al. 1998) was used for head model construction because an individual MRI was not available. Source reconstruction was performed for 567 positions on the cortical surface via Linear Constrained Minimum Variance Beamforming with a regularization parameter of 5% (van Veen et al. 1997). Subsequently, each source was assigned to one of 48 anatomical parcels covering the cortex defined in the automated anatomical atlas (Tzourio-Mazoyer et al. 2002). Within each parcel, sources were aggregated into a single time series by extracting the largest eigenvector. Finally, parcel time courses were jointly orthogonalized for signal leakage reduction (Colclough et al. 2015).

### Data Analysis

Parcel time courses were segmented into 2s epochs with 1s overlap. Each segment was tapered with 7 tapers of the discrete prolate spheroidal sequence for 2Hz spectral smoothing and forwarded to Fourier transformation. Fourier coefficients were averaged over segments, resulting in one power spectrum per parcel and subject. Each power spectrum was corrected for its aperiodic 1/f background estimate using the *fitting oscillations and one over F* (FOOOF) algorithm (Donoghue et al. 2020) in Python 3.9.1. This toolbox additionally allowed for the detection of peaks in the parcels’ power spectra and estimation of peak frequency and peak amplitude. When applying FOOOF, we started with a simple initial *fixed* model on the frequency interval between 3Hz and 48Hz containing only the offset and exponential slope parameters. Next, we evaluated the fit and increased model complexity iteratively until a good fit was achieved for every parcel (Fig. S2 of the Supplementary Material). In this way, we avoided possible pitfalls when using FOOOF (Gerster et al. 2021). The aperiodic 1/f background was subtracted from the original power spectrum. Power values between 4Hz and 30Hz were used for further analysis to reduce the potential influence of low-frequency artifacts and optimize the 1/f-fit.

### Spectral Slowing

The term spectral slowing refers to a shift of spectral energy towards lower frequencies. We investigated spectral slowing by assessing group differences in the power spectrum’s center of mass, referred to as center of energy (CoE) in the following. Power values between 4Hz and 30Hz were considered to reduce the impact of potential low-frequency artifacts. To ensure that all summands in the calculation of the CoE were positive, the spectrum’s minimum was subtracted from each power value before computing as follows:

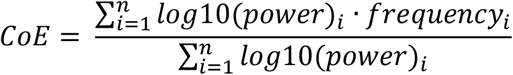

Note that spectral slowing may arise due to different phenomena: 1) A peak shift towards the low frequency range, 2) an increase of peak amplitude in the low frequency range, 3) a decrease of peak amplitude in the high frequency range. In order to disentangle these effects, we first identified a brain region of interest (ROI) based on significant group differences in CoE. Within this ROI, we investigated peak frequency and peak amplitude in detail by means of linear mixed modeling. This analysis was performed once for all peaks detected in the ROI and once using only the largest peak of each parcel. The latter step served to exclude very small, potentially false positive peaks.

### Statistical Analysis

For statistical comparison of CoE, we used cluster-based permutation testing as implemented in Fieldtrip (Maris and Oostenveld 2007). The topographical neighborhood of each parcel was determined by finding all adjacent parcels within 1 cm distance from the parcel border. We report two-sided statistical tests with an alpha level of 5% unless stated otherwise. Parcels showing differences in CoE underwent post-hoc analysis with Linear Mixed Modelling, as implemented in Matlab’s Statistics and Machine Learning Toolbox. We defined separate models to test for group (fixed effect) differences in peak frequency and peak amplitude (dependent variables). Age was included as a covariate together with two-way interactions. Subjects and parcels were defined as random effects. Continuous fixed effects were centered on their mean and APS was chosen as the reference group. Statistical results are reported in the form of *t*-values, indicating the direction of the effect, and corresponding *p*-values.

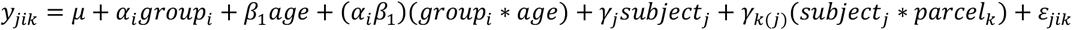

*y*_*jik*_: Response variable; *µ*: Coefficient for the fixed intercept; *α*_i_: Coefficients for group related deviation from the fixed intercept *i* ∈ {1,2}; *β*_1_:Coefficient for covariate age; (*α*_i_*β*_1_): Coefficients for interaction terms *i* ∈ {1,2}; *γ* : Coefficients of subject related random intercept *j* ∈ {1, …, 69}; *γ* : Coefficients of parcel related random intercept nested within subjects *k* ∈ {1, …, 15}; *ε*_i_ ∼*N*(0, *σ*^2^).

## Results

### CBS and PSP vs. HC

We compared resting-state cortical power between CBS patients, PSP patients, and healthy controls. Fig. 1 depicts the group-mean topographies of 1/f-corrected power in the theta (θ: 4Hz–7.5Hz), alpha (α: 8Hz-12.5Hz), low beta (low β: 13Hz-19.5Hz) and high beta band (high β: 20Hz-30Hz). Individual spectra are provided in Fig.S3 of the Supplementary Material. CBS patients and PSP patients showed higher power in the theta and alpha band compared to HC. Theta power was mostly increased in posterior cortical regions, whereas alpha power was increased in central and frontal regions for both CBS (Fig.1A) and PSP patients (Fig.1B).

**Fig. 1:**
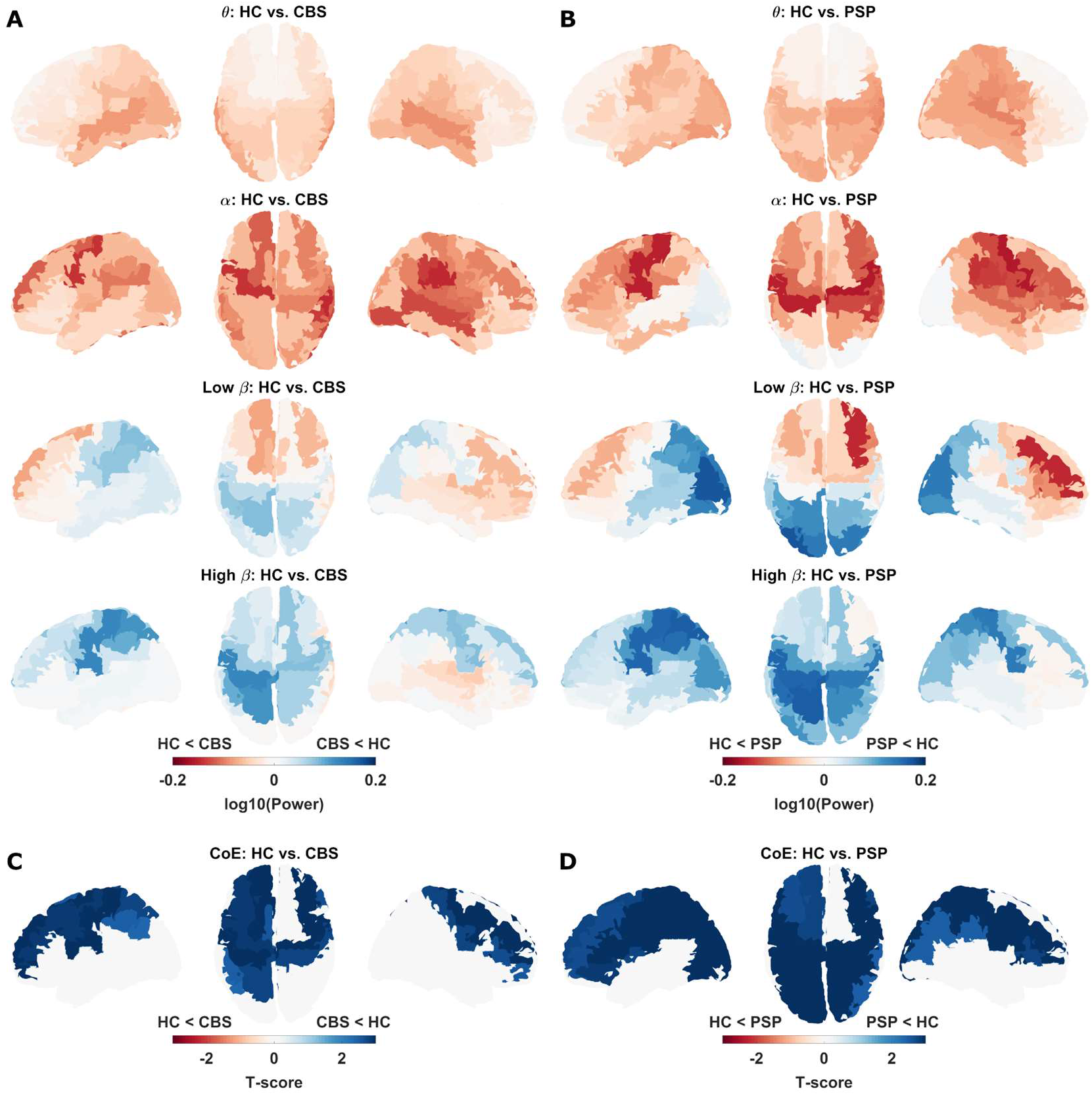
Spectral shift of energy towards lower frequencies in CBS and PSP in comparison to controls. A) Power differences between controls and CBS patients. B) Power differences between controls and PSP patients. C) Statistical comparison of CoE controls vs. CBS patients. D) As C) for controls vs. PSP.

The power differences reversed in sign for the low and the high beta range, indicating larger beta power in controls, particularly in central and parietal cortex. Thus, APS patients exhibited stronger theta/alpha power, but weaker beta power than controls, suggesting a shift of spectral energy towards lower frequencies. The CoE, capturing this shift in a single number, differed from HC for both the CBS and the PSP group in numerous frontal, central and parietal areas (Fig.1C and D, cluster-based permutation test: HC vs. CBS: *p* = 0.004, HC vs. PSP: *p* = 0.002). CBS and PSP, however, did not differ significantly (no clusters found). Therefore, both groups were merged to jointly represent APS in the following. Individual CoE data can be found in Fig. S5A of the Supplementary Material.

### APS vs. PD

When comparing band-limited power between APS and HC, the effects remained significant, as expected (Fig. 2A+C; *p* = 0.001). When comparing band-limited power between APS and PD, we observed a similar pattern as in the comparison to healthy controls. APS had more alpha and less high beta power than PD, although the difference was less strong (Fig. 2B). Again, the CoE was reduced in APS, but the reduction was more local than in the previous comparison, occurring in frontal and central areas exclusively (Fig.2D; *p* = 0.005). These findings demonstrate that APS has a different resting-state oscillatory profile than PD.

**Fig. 2:**
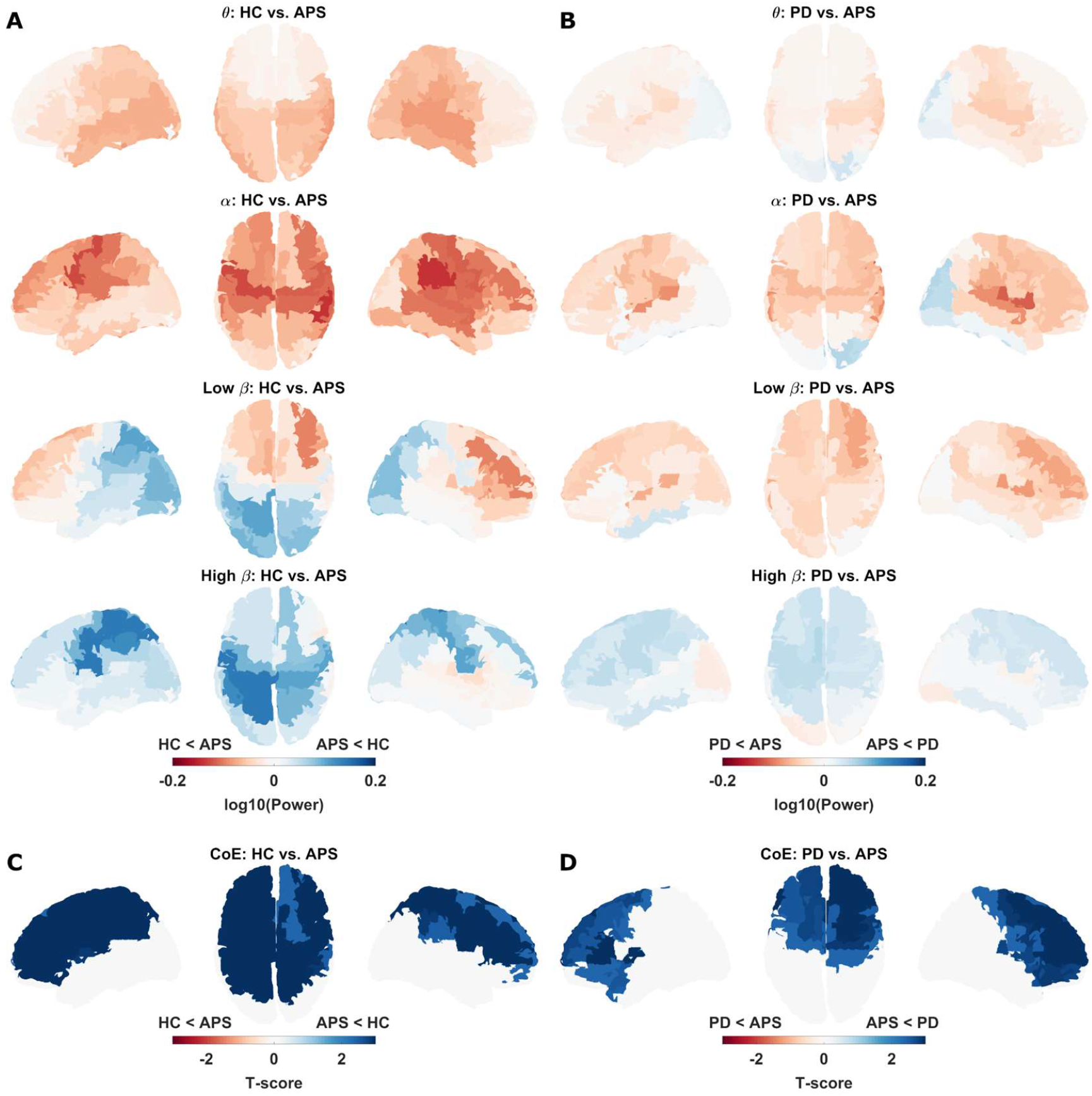
Spectral shift of energy towards lower frequencies in atypical Parkinsonism in comparison to healthy controls and patients with idiopathic Parkinson syndrome. A) APS vs. controls. B) APS vs. PD. C) Statistical comparison of center of energy (CoE) APS vs. controls. D) as C) for APS vs. PD.

The pattern for CBS patients did not change qualitatively when mirroring brain images such that the hemisphere contralateral to the symptom-dominant bodyside was always right (Fig. S4 of the Supplementary Material). The average CoE within the brain region with a significant difference to controls did not correlate with UPDRS-III scores (r_Spearman_ = 0.151, *p* = 0.622), MoCA scores (r_Spearman_ = 0.209, *p* = 0.514), TULIA scores (r_Spearman_ = 0.107, *p* = 0.727), or Goldenberg scores (r_Spearman_ = 0.220, *p* = 0.470) in the CBS cohort.

### Spectral Peaks

The CoE analysis revealed a shift of energy towards lower frequencies in APS patients. Next, we describe this shift in more detail through an analysis of spectral peaks. A shift of spectral power may arise due to changes in peak frequency, peak amplitude, or both. We sought to disentangle these effects by means of linear mixed modeling. This analysis was performed for a region of interest (ROI) showing significant differences in CoE in the comparisons APS vs. HC and APS vs. PD (Fig. 3A).

**Fig. 3:**
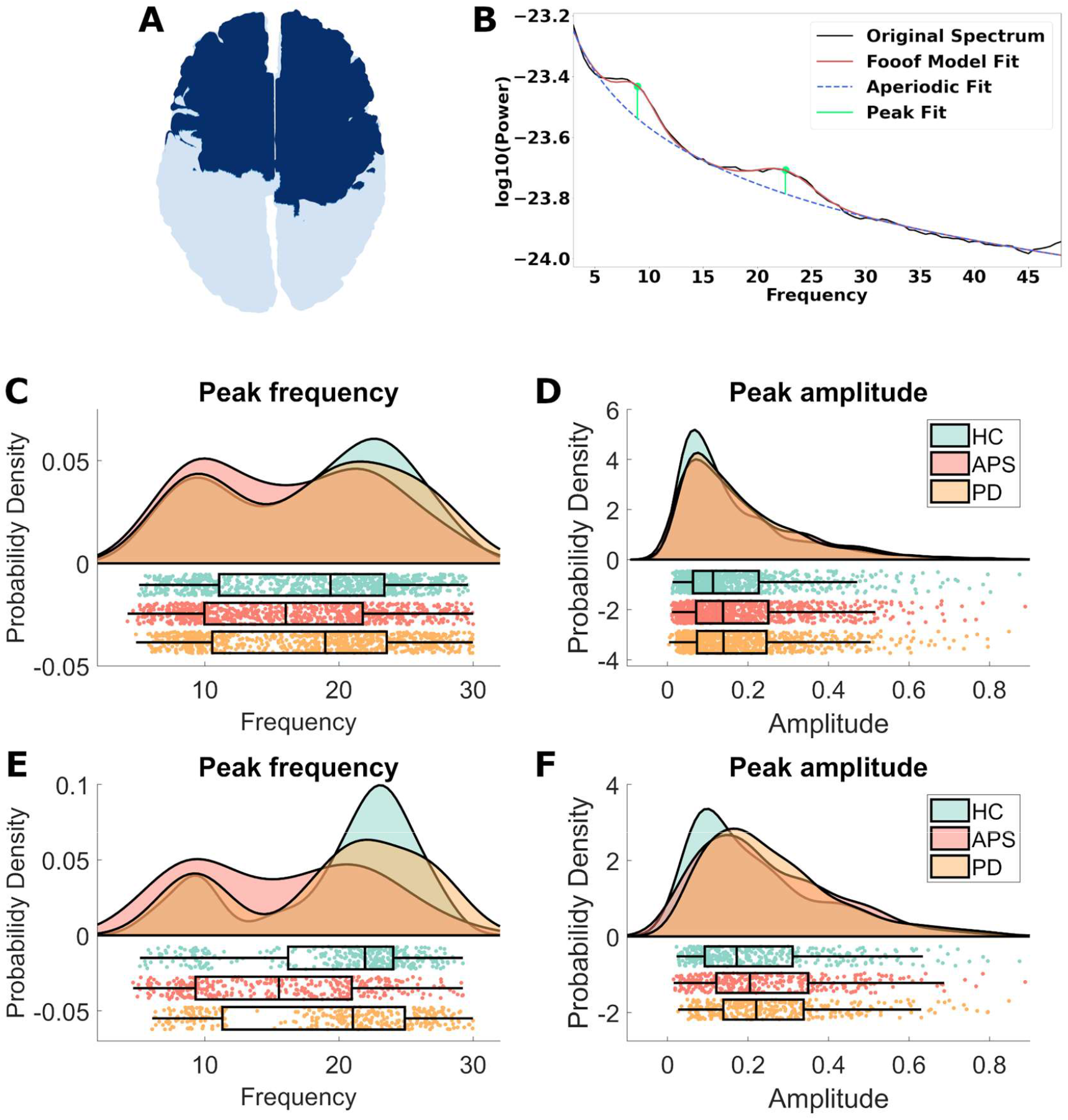
Peak frequency and peak amplitude distribution for APS, PD and HC. A) ROI for analysis of peaks. B) Example of parcel spectrum (black) and FOOOF estimate of the aperiodic background (blue dotted line) and the periodic components (red) with individual peak fits (peak 1: frequency = 8.937, amplitude = 0.108, peak 2: frequency = 22.599, amplitude = 0.078). C) Probability density of peak frequencies. All peaks found in the ROI considered. D) Probability density of peak amplitude. All peaks found in the ROI considered. E) as C), only the largest peaks per parcel considered. F) as D), only the largest peak per parcel considered. ROI: Region of interest, HC: Healthy controls, APS: Atypical Parkinson syndromes, PD: Parkinson’s disease.

Figure 3C illustrates the distribution of peak frequencies in the ROI for APS patients, PD patients and HC. Peak frequencies were extracted with the FOOOF toolbox (see Fig.4B for an example). The distribution was bimodal for all groups, with peak frequencies clustering around 9Hz and 22Hz. For this reason, we performed separate statistical comparisons for the theta/alpha (4Hz-13Hz) and the beta band (13Hz-30Hz). In Fig. S5B of the Supplementary Material, we present an alternative analysis based on the entire frequency range (4Hz-30Hz).

**Fig. 4:**
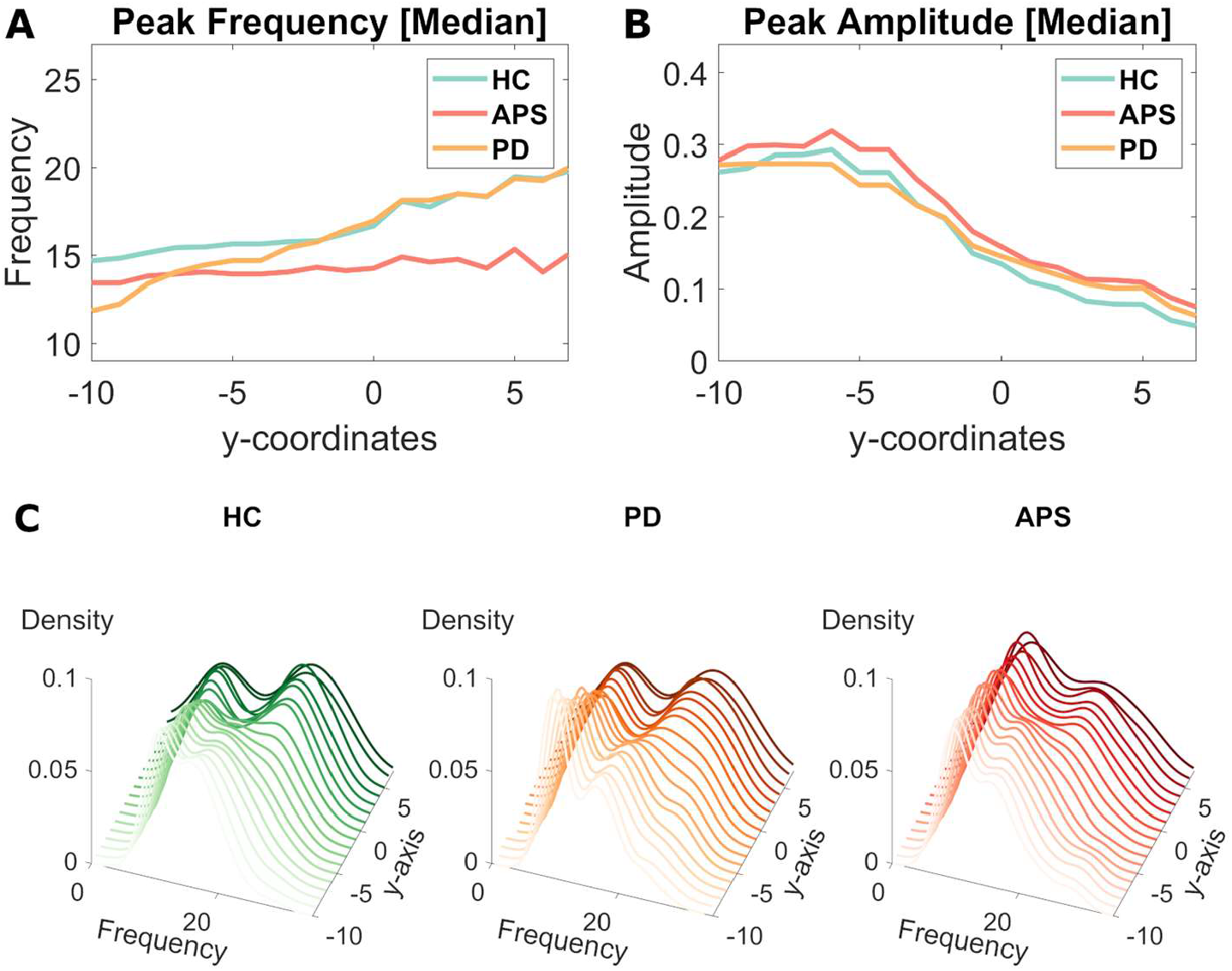
Patients with atypical Parkinson syndromes lack anterior-posterior gradient in peak frequency. A) Median peak frequency along the posterior to anterior axis (y = 0 at anterior commissure). B) Median peak amplitude along the posterior to anterior axis. C) Probability density of peak frequencies for eighteen coronal slices along the anterior-posterior axis (slice thickness: 5 cm, step-size: 1 cm). HC: Healthy controls, APS: Atypical Parkinson Syndrome, PD: Parkinson’s disease.

### Theta/alpha oscillations

#### Peak frequency

There was no significant difference in theta/alpha peak frequency between the groups, neither when considering all peaks found in the ROI (*t*_*HC*_ = 0.011, *p*_*HC*_ = 0.992; *t*_*PD*_ = -0.154, *p*_*PD*_ = 0.877; *t*_*age*_ = 0.545, *p*_*age*_ = 0.586), nor when considering only the highest peak per ROI parcel (*t*_*HC*_ = 0.288, *p*_*HC*_ = 0.773; *t*_*PD*_ = 0.007, *p*_*PD*_ = 0.995; *t*_*age*_ = 0.669, *p*_*age*_ = 0.504).

#### Peak amplitude

Theta/alpha peak amplitudes were increased in APS patients in comparison to controls (*t* = -2.556, *p* = 0.011), but not in comparison to PD (*t* = -0.684, *p* = 0.494). Age did not influence theta/alpha peak amplitude (*t* = -1.074, *p* = 0.283). Similar results were obtained when considering only the largest peaks per parcel (*t*_*HC*_ = -2.639, *p*_*HC*_ = 0.009; *t*_*PD*_ = -0.844, *p*_*PD*_ = 0.399; *t*_*age*_ = -0.946, *p*_*age*_ = 0.345). We note that the amplitude effect in the frontal ROI was most likely due to differences in the alpha band (Fig. S6 of the Supplementary Material). Theta power differences occurred in more posterior areas.

### Beta oscillations

#### Peak frequency

Beta peak frequencies were lower in APS patients compared to PD patients (*t* = 2.799, *p* = 0.005) and controls (*t* = 2.355, *p* = 0.019) when considering all peaks found in the ROI (Fig. 3C). Across the groups, peak frequency did not depend on age (*t* = 0.78, *p* = 0.435), but peak frequency decreased with age in the PD group (*t* = -2.313, *p* = 0.021) and healthy controls specifically (*t* = -1.982, *p* = 0.048). The reduction of beta peak frequency remained significant when considering the largest beta peaks instead of all beta peaks (*t*_*PD*_ = 2.981, *p*_*PD*_ = 0.003; *t*_*HC*_ = 2.749, *p*_*HC*_ = 0.006).

#### Peak amplitude

Beta amplitudes were similar between groups when considering all peaks (*t*_*HC*_ = 0.025, *p*_*HC*_ = 0.98; *t*_*PD*_ = 0.212, *p*_*PD*_ = 0.832). The main effect of age was not significant (*t* = -0.753, *p* = 0.451) and there were no interactions (all |*t-*values| < 1.07 and *p-*values > 0.285). Amplitude differences remained insignificant when considering the largest beta peaks (*t*_*HC*_ = 0.073, *p*_*HC*_ = 0.942; *t*_*PD*_ = 0.553, *p*_*PD*_ = 0.581).

These results are evidence of a genuine slowing of fronto-central beta oscillations in APS, in the sense of a peak shift without changes in amplitude. We note that the linear mixed modeling accounted for the fact that each participant contributed a different number of peaks. Thus, our findings cannot be explained by APS patients having less beta peaks overall.

#### The topography of slowing

We investigated the topography of spectral slowing in APS by analyzing the anterior-posterior gradient in peak frequency. APS patients lacked the growing predominance of beta over alpha peak frequencies when moving from the posterior to the anterior end of the brain. This gradient was visible in the PD and HC group (Fig.5A and 5C). Peak amplitude did not show this gradient after 1/f correction.

## Discussion

In this study, we show to our knowledge for the first time, in a comparably large cohort, that resting-state oscillatory activity differs between APS patients, PD patients and HC. APS patients exhibit a shift of spectral energy towards lower frequencies, resulting from increased amplitude of theta and alpha oscillations and a slowing of frontal and central beta oscillations independent of amplitude. Further, oscillations in APS do not positively scale in peak frequency along the posterior-to-anterior axis, as they do in PD patients and healthy participants. Thus, APS has a characteristic spatio-spectral signature, implying that APS and IPS might be distinguishable based on MEG, and possibly EEG, recordings.

### Previous studies

Slowing of neural oscillations has rarely been investigated in the context of APS. To our knowledge, only Montplaisir and colleagues reported enhanced sensor-level delta and theta power in relation to alpha and beta power for six PSP patients contrasted to six healthy controls (Montplaisir et al. 1997). Here, we corroborate and extend these findings by demonstrating that slowing is a prominent feature of CBS and PSP patients.

### Spectral slowing in other diseases

The changes in power reported here are reminiscent of pathological alterations observed in other neurological diseases such as multiple sclerosis (van Schependom et al. 2021), hepatic encephalopathy (Butz et al. 2013), and AD. Especially for AD, numerous studies have reported spectral slowing (Fernández et al. 2006; Wiesman et al. 2022), increased theta activity and decreased alpha and beta activity (Huang et al. 2000; Berendse et al. 2000; Fernández et al. 2002; Osipova et al. 2005; Fernández et al. 2006; Dauwels et al. 2011; Haan et al. 2008; Jelic et al. 2000; Coben et al. 1985; Jafari et al. 2020; Nakamura et al. 2018; Wiesman et al. 2022). These spectral changes have been linked to disease progression (Jelic et al. 2000; Coben et al. 1985; Huang et al. 2000; Babiloni et al. 2009; Nakamura et al. 2018) and were successfully used to differentiate AD patients from healthy controls (Gouw et al. 2021).

Because CBS and PSP, like AD, are tauopathies exhibiting spectral slowing, it is conceivable that comparable neurodegenerative processes lead to a common spectral pattern, characterized by a loss of energy in the high-beta band and increased power in the theta band. This neurodegenerative process might culminate in a complete decay of beta and alpha oscillations. The spectral differences between APS and PD observed here might result from the relationship between spectral slowing and cortical atrophy. PD is characterized by local midbrain degeneration (Lang and Lozano 1998) rather than severe cortical degeneration. Although several gray matter changes are detectable in the neocortex of PD patients (Ramírez-Ruiz et al. 2005; Nishio et al. 2010; Lee et al. 2013), frontal atrophy is not as prominent as in CBS patients (Josephs et al. 2008; Whitwell et al. 2010; Lee et al. 2011; Josephs et al. 2010; Albrecht et al. 2017; Dutt et al. 2016), or PSP patients (Paviour et al. 2006a, 2006b; Cordato et al. 2002; Gröschel et al. 2004; Stamelou et al. 2011; Josephs et al. 2008; Brenneis et al. 2004; Whitwell et al. 2011; Schofield et al. 2011).

### Tauopathy and neuronal oscillations

Several studies support a direct link between tauopathy, neurodegeneration and neuronal oscillations. Atrophy patterns and pathological tau burden tend to colocalize in AD (Xia et al. 2017; La Joie et al. 2020; Mak et al. 2018), CBS (Dickson et al. 2002; McMillan et al. 2016), and PSP (Nicastro et al. 2020). Although tau burden does not necessarily correlate with the current severity of atrophy in PSP patients (Schofield et al. 2011), it was found to be highly predictive of future atrophies in AD (La Joie et al. 2020).

The link between tau pathology and atrophy is relevant for the current study as previous studies suggested both tau-burden (Stomrud et al. 2010; Smailovic et al. 2018; Coomans et al. 2021) and atrophy (Nakamura et al. 2018; Fernández et al. 2003; Grunwald et al. 2007; Briels et al. 2020) to modulate spectral properties. Tau-related cerebrospinal fluid biomarkers have been found to be associated with elevated theta power in elderly, healthy participants (Stomrud et al. 2010) and reduced alpha and beta power in AD patients (Smailovic et al. 2018). Furthermore, tau accumulation was related to posterior slowing of oscillatory activity in a recent MEG study (Coomans et al. 2021) and human-tau-transgenic mice exhibit power increases in the delta and theta band in parietal areas reminiscent of spectral changes observed in AD patients (Das et al. 2018). Oscillatory markers were also found to correlate with amyloid-beta accumulation in AD (Nakamura et al. 2018; Wiesman et al. 2022), a pathological process assumed to precede tau pathology (Buchhave et al. 2012; Jack et al. 2013), suggesting general sensitivity of oscillatory metrics to proteinopathy (Wiesman et al. 2022).

Whether the observations linking tau, atrophy and oscillations are transferable from AD to APS is unclear. Here, we found that APS, like AD, is associated with spectral slowing. Interestingly, the topography of slowing seems to be disease-specific, with slowing predominating in areas most affected by cortical neurodegeneration. In APS, slowing occurs in frontal, rolandic and parietal areas, matching the reported atrophy patterns (Boeve et al. 1999; Josephs et al. 2008; Whitwell et al. 2010) and topography of tau accumulation (Forman et al. 2002; Cho et al. 2017; McMillan et al. 2016) in CBS and, and with less cortical involvement, in PSP (Nicastro et al. 2020). These results suggest that oscillatory slowing could potentially serve as a marker of cortical neurodegeneration in the future, complementing structural and molecular imaging techniques.

### Peak frequency gradient

In keeping with our results, a recent MEG study in young, healthy adults found beta peak frequency to scale positively along the posterior-to-anterior axis (Mahjoory et al. 2020). This pattern appears to be absent in APS patients, who lacked the transition from alpha to beta dominance upon reaching frontal cortex. Thus, the gradient itself might carry information relevant to diagnosis. In contrast to the report by Mahjoory and colleagues., we did not find theta oscillations to dominate in frontal cortex. This might be due to age differences between the studies. Here, we included elderly participants, whereas Mahjoory and colleagues recruited from a student population. Theta power is known to decrease in healthy aging (Vlahou et al. 2014). In our sample, we observed an age-related decrease of beta peak frequency in the PD and HC group, next to a much stronger disease-related shift in APS patients, independent of age. This suggests that age-related peak frequency shifts (Rossiter et al. 2014; Barry and Blasio 2017) can be modulated by disease. Both, age- and disease-related spectral slowing, of course, might be caused by neurodegeneration, unfolding at different time scales in healthy aging and disease.

### Common patterns in APS and PD

An increase of low alpha/theta peak amplitude relative to controls was observed in both APS and PD. This finding aligns with previous studies reporting increased theta and low alpha power in PD in comparison to controls (Bosboom et al. 2006; Stoffers et al. 2007) and further suggests that parkinsonian syndromes have both common and distinct alterations in cortical power. The similarities in low oscillatory activity may emerge due to common pathological subcortical changes, e.g. in the locus coeruleus, that is involved in PD (Dickson 2012), PSP (Dickson 2012; Dickson 1999) and CBS (Dickson et al. 2002; Dickson 1999) and has been implicated in oscillatory slowing in rats (Berridge et al. 1993). While the increase in theta power can be considered a common aspect of slowing in Parkinsonism, the shift of beta peak frequency in frontocentral areas seems to be specific to APS.

### Slowing and symptom severity

In this study, we found no indication that spectral slowing relates to CBS symptom severity when correlating the ROI-mean CoE to UPDRS, MoCA, TULIA or Goldenberg scores, indicating that slowing reflects disease entity or, possibly, severity of atrophy rather than the current symptoms. As reported in a previous study in PSP patients, the severity of atrophy does not necessarily reflect symptom severity (Schofield et al. 2011). This finding might be transferable to spectral slowing. A definite answer to this question, however, requires larger samples. The CBS group studied here contained only 13 patients and was heterogeneous with respect to clinical symptoms.

### Distinguishing atypical parkinsonian syndromes

While we found differences between APS and PD patients, we did not find differences between the APS subgroups CBS and PSP. Niethammer and colleagues have made considerable progress in this direction by successfully distinguishing CBS from PSP with positron emission tomography (Niethammer et al. 2014). Clearly, it would be desirable to achieve this with electrophysiological techniques as well, which do not require injection of radioactive tracers and, in case of EEG, are widely available. Our results set the ground for future research in this direction, highlighting the importance of the distribution of peak frequencies rather than power in a priori defined frequency bands alone. In fact, this concept might be applicable to a wider range of M/EEG research topics. Distinguishing atypical parkinsonian syndromes might require inclusion of more patients, improving subgroup homogeneity and/or considering further features of brain activity, such as connectivity. In this way, it might be possible to classify individual patients in the future.

## Conclusions

Here, we demonstrate that CBS and PSP have a characteristic spatio-spectral resting-state profile distinguishing them from PD and HC. These findings contribute to the understanding of how neurodegeneration affects neuronal oscillations and might be of diagnostic value in the future.

## Acknowledgements

This study was founded by the Else Kröner Fresenius Stiftung. We would like to thank Lucie Winkler and Julian Caspers for their help with neuropsychological scores and imaging.

