## Supplementary material for "Slowing of Frontocentral Beta Oscillations in Atypical Parkinsonism": Tab. S1; Fig. S1; Fig. S2; Fig. S3; Fig. S4; Fig. S5; Fig. S6

Marius Krösche[1], Silja Kannenberg[1], Markus Butz[1],

Christian J. Hartmann[1,2], Esther Florin[1], Alfons Schnitzler[1,2], Jan Hirschmann[1]

**Demographics and test battery**

Demographic and disease-related information on patient groups are displayed in Table S1. UPDRS-III motor scores were available for the CBS and the PD group. CBS patients underwent a neuropsychological test battery to evaluate the severity of apraxia (Goldenberg and TULIA) and cognitive impairment (MoCA). Laterality was defined as right minus left hemibody scores. Relative laterality scores are the average of UPDRS III, Goldenberg and TULIA lateralities, each normalized by the highest possible difference, and reflect overall lateralization of symptoms. Cbs11 did not receive a MoCA score due to severe aphasia. For Cbs04, Cbs05 and Cbs13 the sub-items on visuospatial abilities were disregarded due to severe motor impairments that hampered drawing abilities. For PD14, two sub-items of the UPDRS-III were missing.

*Tab. S1: Demographic and diagnostic information on APS patients and PD patients. CBS: Corticobasal syndrome; PSP: Progressive supranuclear palsy syndromes; UPDRS: Unified Parkinson's disease rating scale; Goldenberg: Goldenberg's apraxia test; TULIA: Test of upper limb apraxia; MoCA: Montreal Cognitive Assessment.*

| Patient | Age | Gender | Diagnosis (possible/ probable) | UPDRS III (sum) | UPDRS III Laterality | Goldenberg (sum) | Goldenberg Laterality | TULIA (sum) | TULIA Laterality | Relative Laterality | MoCA (sum) |
|---|---|---|---|---|---|---|---|---|---|---|---|
| **CBS01** | 57 | f | possible | 38 | -6 | 77 | -3 | 22 | -2 | -0.13 | 27 |
| **CBS02** | 61 | f | probable | 58 | 11 | 33 | 25 | 12 | 4 | 0.4 | 9 |
| **CBS03** | 76 | f | possible | 47 | 4 | 67 | -1 | 21 | 1 | 0.05 | 16 |
| **CBS04** | 76 | f | probable | 60 | 1 | 4 | 0 | 2 | 2 | 0.06 | 17 |
| **CBS05** | 60 | f | probable | 41 | -2 | 10 | -6 | 8 | -2 | -0.12 | 5 |
| **CBS06** | 61 | m | probable | 47 | -15 | 78 | -2 | 23 | 1 | -0.10 | 26 |
| **CBS07** | 60 | m | possible | 76 | -4 | 64 | -8 | 18 | -4 | -0.21 | 17 |
| **CBS08** | 69 | f | probable | 83 | 8 | 50 | 6 | 17 | -3 | 0.03 | 11 |
| **CBS09** | 52 | f | possible | 11 | 4 | 80 | 0 | 24 | 0 | 0.03 | 19 |
| **CBS10** | 65 | m | probable | 20 | 12 | 52 | 20 | 17 | -1 | 0.23 | 29 |
| **CBS11** | 72 | f | probable | 78 | -5 | 23 | -15 | 5 | -3 | -0.25 | - |
| **CBS12** | 52 | f | probable | 16 | -6 | 58 | -18 | 22 | -2 | -0.25 | 14 |
| **CBS13** | 71 | m | probable | 55 | 2 | 4 | 0 | 3 | 1 | 0.04 | 13 |
| **PSP01** | 68 | f | probable | | | | | | | | |
| **PSP02** | 64 | f | probable | | | | | | | | |
| **PSP03** | 73 | m | probable | | | | | | | | |
| **PSP04** | 71 | f | probable | | | | | | | | |
| **PSP05** | 70 | m | probable | | | | | | | | |
| **PSP06** | 67 | m | probable | | | | | | | | |
| **PSP07** | 71 | m | probable | | | | | | | | |
| **PSP08** | 79 | m | probable | | | | | | | | |
| **PSP09** | 78 | f | probable | | | | | | | | |
| **PSP10** | 74 | f | possible | | | | | | | | |
| **PD01** | 47 | m | | 62 | -8 | | | | | | |
| **PD02** | 68 | m | | 33 | 3 | | | | | | |
| **PD03** | 71 | m | | 50 | 0 | | | | | | |
| **PD04** | 52 | m | | 47 | -4 | | | | | | |
| **PD05** | 68 | m | | 26 | 5 | | | | | | |
| **PD06** | 74 | m | | 37 | 17 | | | | | | |
| **PD07** | 71 | m | | 24 | -9 | | | | | | |
| **PD08** | 58 | f | | 29 | -5 | | | | | | |
| **PD09** | 66 | m | | 22 | -4 | | | | | | |
| **PD10** | 58 | m | | 7 | -7 | | | | | | |
| **PD11** | 58 | f | | 31 | 2 | | | | | | |
| **PD12** | 79 | m | | 41 | 1 | | | | | | |
| **PD13** | 65 | f | | 40 | -4 | | | | | | |
| **PD14** | 58 | m | | 41 | 1 | | | | | | |
| **PD15** | 67 | m | | 28 | 4 | | | | | | |
| **PD16** | 65 | f | | 32 | 3 | | | | | | |
| **PD17** | 69 | m | | 42 | 1 | | | | | | |
| **PD18** | 73 | m | | | | | | | | | |
| **PD19** | 68 | m | | 38 | -1 | | | | | | |
| **PD20** | 64 | m | | 43 | -1 | | | | | | |
| **PD21** | 55 | m | | 33 | 0 | | | | | | |
| **PD22** | 54 | m | | 40 | -6 | | | | | | |
| **PD23** | 58 | m | | 53 | -3 | | | | | | |

**Cortical parcellation**

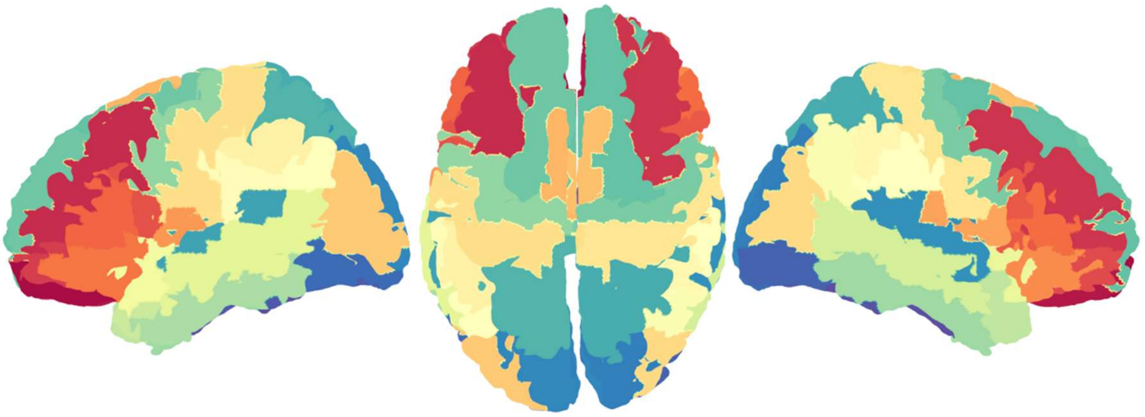

*Fig. S1.: Segmentation of cortex into 48 anatomical regions based on the AAL atlas. Rolandic operculum L/R, postcentral gyrus L/R, precentral gyrus L/R; Parietal superior cortex L/R, parietal inferior cortex L/R, supramarginal gyrus L/R, angular gyrus L/R; Superior frontal gyrus L/R, middle frontal gyrus L/R, inferior frontal gyrus pars orbitalis L/R, superior frontal gyrus pars orbitalis L/R, middle frontal gyrus pars orbitalis L/R, inferior frontal gyrus pars opercularis L/R, inferior frontal gyrus pars triangularis L/R, supplementary motor area L/R; Superior temporal gyrus L/R, superior temporal pole L/R, middle temporal gyrus L/R, middle temporal pole L/R, inferior temporal gyrus L/R; Superior occipital gyrus L/R, middle occipital gyrus L/R, inferior occipital gyrus L/R; Cerebellum L/R (not displayed).*

**Fitting aperiodic and periodic spectral components**

For inter-subject comparison, we normalized power spectra by subtracting their 1/f aperiodic component computed with the toolbox *fitting oscillations & one over f* by Donoghue and colleagues (Donoghue et al. 2020). FOOOF also facilitated the estimation of peak frequency and peak amplitude of oscillations. The model estimate was fitted on the 3Hz to 48Hz interval to ensure the best possible fit on the analysis interval between 4Hz and 30Hz. Parameter settings were adapted after visual inspection if needed to account for possible pitfalls when using FOOOF (Gerster et al. 2021).

Four different model fits with different flexibility were used. 1. *Fixed model* fitting with standard setting *'fixed'*, based on estimation of two parameters for offset and slope of the aperiodic fit. 2. *Knee model* fitting with setting *'knee'*, based on estimation of the offset, slope and a third parameter adding flexibility to the slope. 3. *Offset model* fitting the standard *fixed model* but shifting the offset between 2Hz and 4Hz to improve the aperiodic fit. 4. *Stepwise model* fitting a first *fixed model* from 3Hz to a border-value and a second *fixed model* from the border-value to 48Hz. Examples are given below. We note that many fits would have been of insufficient quality if the same model had been applied in all cases.

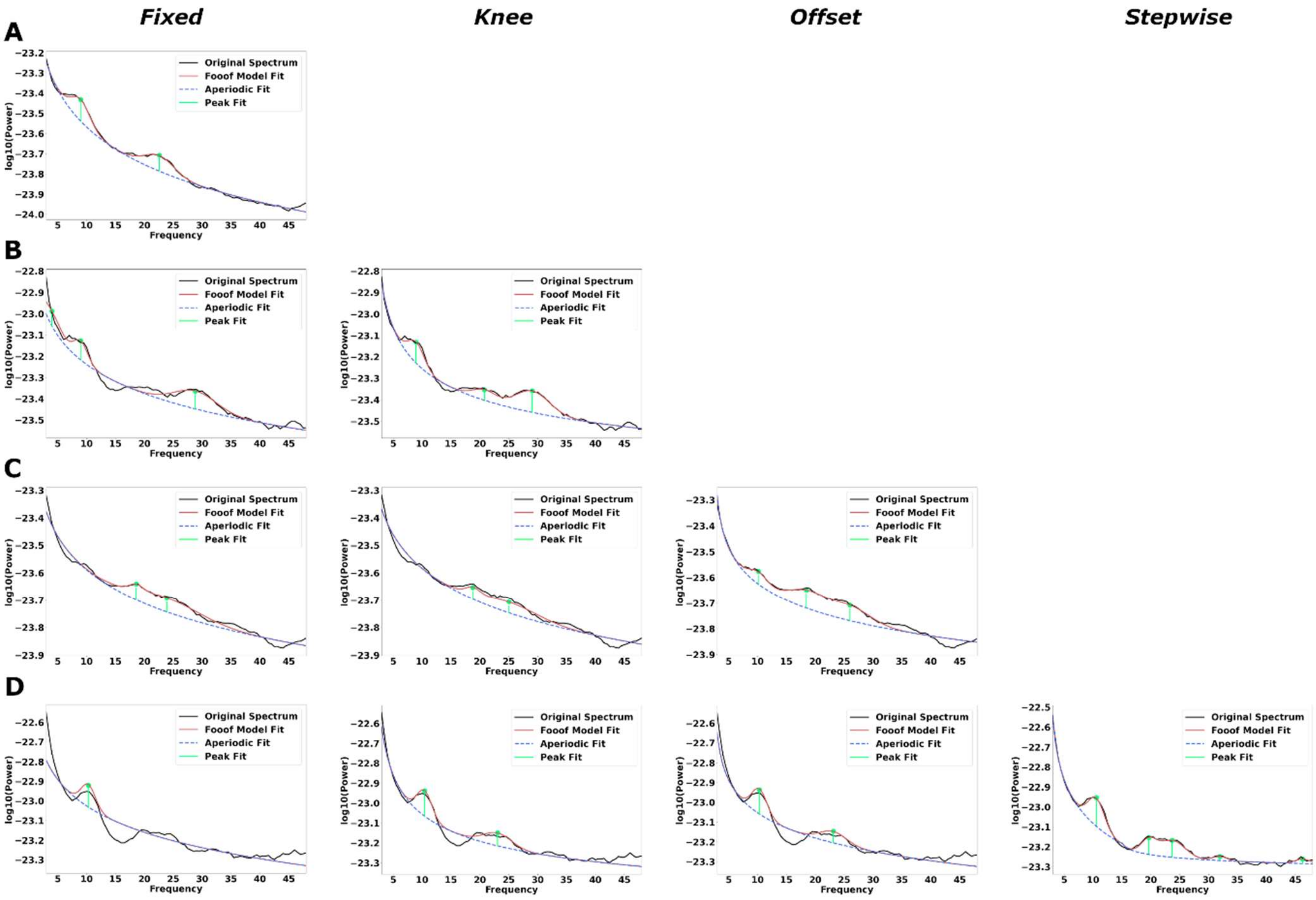


*Fig. S2: Example spectra for different model fits. A) Fixed model: Successful fit with two peak estimates. B) Fixed model: Unsuccessful fit. Offset estimate far from original spectrum, non-existing theta oscillation added, low-beta oscillation not detected, and aperiodic fit too high to avoid trough at ~14Hz. Knee model: Successful fit with three peak estimates. C) Fixed model:  Unsuccessful fit. Offset estimate far from original spectrum, low alpha oscillation not detected. Knee model: Unsuccessful fit. Similar to fixed model. Offset model: Successful fit. D) Fixed model: Unsuccessful fit. Offset estimate far from original spectrum, aperiodic fit too high to avoid trough at ~15Hz, does not capture beta activity. Knee model: Unsuccessful fit. Aperiodic fit too high to avoid trough at ~15Hz. Offset model: Unsuccessful fit. No improvement compared to knee model. Stepwise model: 1.Fixed model from 3Hz to 15Hz. 2.Fixed model from 15Hz to 48Hz. The joint model captures oscillatory activity correctly.*

**Individual spectra within region of interest**

Individual spectra for all parcels and all participants are displayed separately for each group in Fig. S3. The APS and the PD cohort both showed higher power than controls  in the in the theta and alpha band. Differences between APS and PD emerged in the beta band. PD, but not APS, spectra had a gap around 15 Hz. The absence of this gap reflects the beta peak shift in APS.

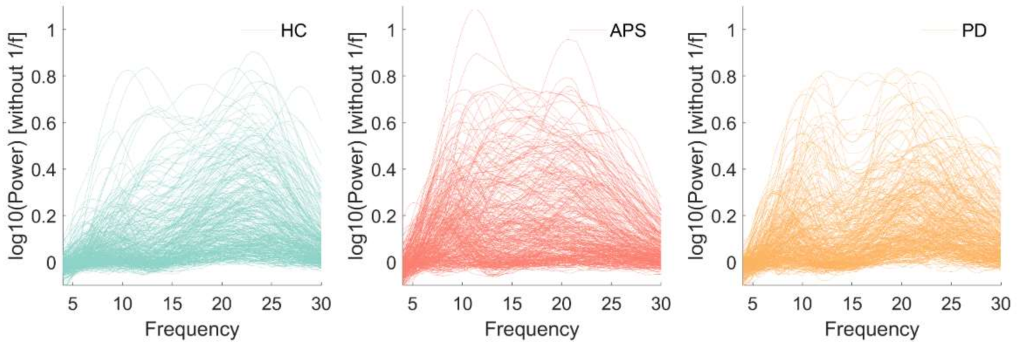


*Fig. S3: Power spectra of all cortical parcels. HC: Healthy controls, APS: Atypical parkinsonian syndromes, PD: Parkinson´s disease.*

**Accounting for symptom asymmetry**

Parkinsonism, and CBS in particular, is usually lateralized, although symptoms and atrophy patterns are symmetric in some patients (Hassan et al. 2010; Armstrong et al. 2013) and lateralization was mild in our sample (Tab. S1). In order to test the influence of asymmetry, we mirrored power topographies of the CBS group such that the hemisphere contralateral to the more affected body side always ended up on the right. This step hardly affected the results (Fig. S4), suggesting at most a minor influence of lateralization.

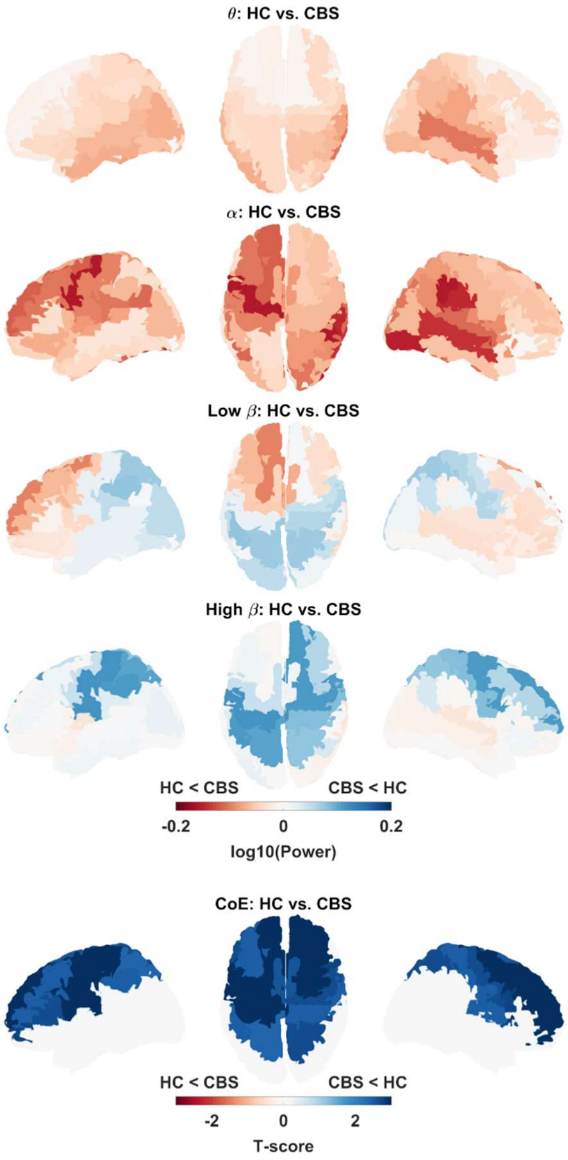


*Fig. S4: Power and CoE difference between CBS and HC after left-right mirroring according to the relative laterality scores (see Tab. S2).*

**Spectral slowing in individuals**

Fig. S5A displays the ROI-average CoE for each participant. Fig. S5B shows the individual number of high-beta peaks relative to the total number of peaks in frequency range from 4 to 30 Hz. This was diminished in APS patients compared to PD patients. A general linear model with the factor disease and the covariate age revealed a group effect. APS patients had a lower proportion of high-beta peaks than PD ($t$ = 2.2768, $p$ = 0.0262) and HC ($t$ = 3.1161, $p$ = 0.0028), consistent with the results presented in the main paper.

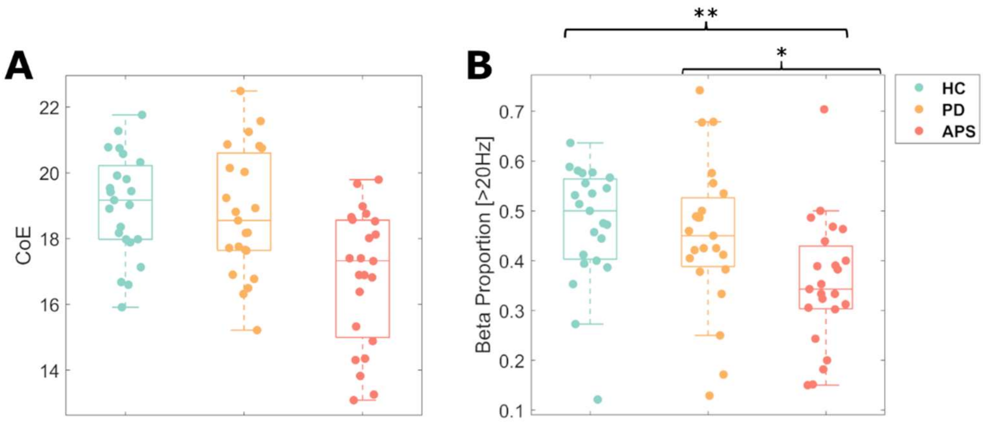


Fig. S5: Individual data A). ROI-average center of energy values for each participant. B) Proportion of high-beta peaks for each participant. HC: Healthy controls (green), PD: Idiopathic Parkinson syndrome (orange), APS: Atypical parkinsonian syndrome (red).

**Whole-brain differences in theta and alpha power**

To understand how the difference in theta/alpha peak amplitude detected in the ROI (compare Fig. 3, main paper) relates to whole-brain topographies of theta and alpha power differences, we performed whole-brain, cluster-based permutation tests (Fig. S6). Alpha power differences between APS and controls localized to bilateral frontal cortex, roughly consistent with the ROI used in the main paper ($p$ = 0.0106). Theta power differences between APS and controls ($p$ = 0.0017) as well as between PD and controls ($p$ = 0.0056) were observed in more posterior regions. These results indicate that the peak amplitude differences reported in the main paper are mainly driven by alpha oscillations.

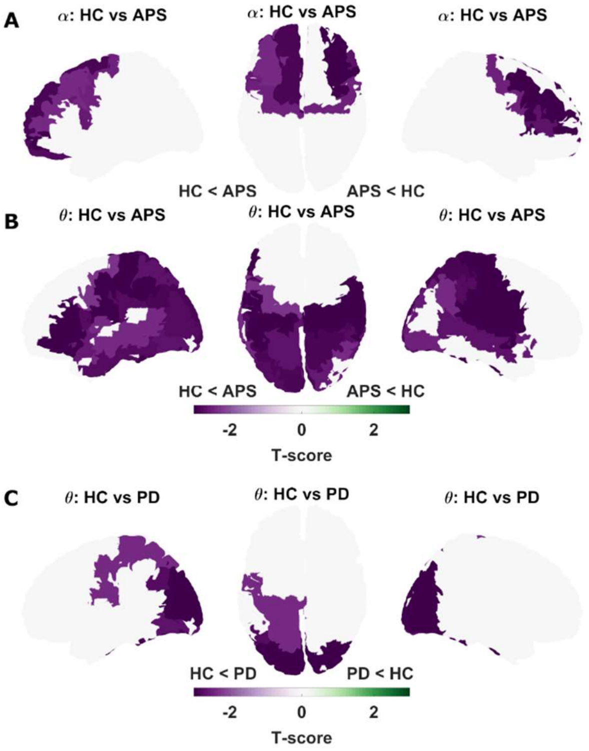


*Fig. S6: Whole-brain power differences. A. Alpha band power difference between APS and HC. B. Theta band power difference between APS and HC. C. Theta band power difference between PD and HC.*
